# Risk factors associated with Parkinson’s disease: An 11-year population-based South Korean study

**DOI:** 10.1101/253690

**Authors:** Hyoung Seop Kim, Hong-Jae Lee, Jiook Cha, Junbeom Kwon, Hyunsun Lim

## Abstract

**Objective:** To validate various known risk factors of Parkinsonism and to establish basic information to formulate public health policy by using a 10-year follow-up cohort model.

**Methods:** This population based nation-wide study was performed using the National Health Insurance Database of reimbursement claims of the Health Insurance Review and Assessment Service of South Korea data on regular health check-ups in 2003 and 2004, with 10 years’ follow-up.

**Results:** We identified 7,746 patients with Parkinsonism. Old age, hypertension, diabetes, depression, anxiety, taking statin medication, high body mass index, non-smoking, non-alcohol drinking, and low socioeconomic status were each associated with an increase in the risk of Parkinsonism (fully adjusted Cox proportional hazards model: hazard ratio (HR) 1.259, 95% confidence interval (CI) 1.194–1.328 for hypertension, HR 1.255, 95% CI 1.186–1.329 for diabetes, HR 1.554, 95% CI 1.664–1.965 for depression, HR 1.808, 95% CI 1.462–1.652 for anxiety, and HR 1.157, 95% CI 1.072–1.250 for taking statin medication).

**Conclusions:** In our study, old age, depression, anxiety, and a non-smoker status were found to be risk factors of Parkinsonism, in agreement with previous studies. However, sex, hypertension, diabetes, taking statin medication, non-drinking of Alcohol, and lower socioeconomic status have not been described as risk factors in previous studies and need further verification in future studies.

---

Parkinsonism is one of the most common neurodegenerative diseases, affecting > 1% of the elderly population.^1^ With the marked increase in the mean age of the global population, the prevalence and incidence of Parkinsonism have also increased.^2^ The impact of Parkinsonism on society is a major concern worldwide, including in South Korea. Many genetic risk factors have been discovered in recent years,^3^ but these account for only a minor portion of Parkinsonism, as the disease etiology involves an interplay of both genetic and environmental factors. Most of these factors remain unknown; thus, investigating the distribution and characteristics of the condition is important for identifying further etiologic factors and planning public health policies.^4^

In South Korean, medical care is delivered via the National Health Insurance Service (NHIS) in which most people are obligated to enroll. All medical health insurance-related data are collected into a central database. Additionally, South Koreans receive regular health checkups with the support of the National Health Insurance Corporation (NHIC) after the age of 40 years. Many studies have investigated the relationship between Parkinsonism and some probable risk factors.^5-8^ These include increasing age,^5^ alcohol-use disorder,^6^ urbanization and exposure to pesticides,^9^ cardiovascular changes,^7^ depression^10^, and anxiety,^11^ while smoking has been proven to a preventative factor against development of Parkinsonism.^12, 13^ However, the relationship between socioeconomic status and the onset of Parkinsonism is unknown.

We hypothesized that the relationship of known risk factors and socioeconomic status with the onset of Parkinsonism could be determined using data from both the regular health checkup and health insurance databases of South Korea. We therefore sought to validate the role of various known risk factors for Parkinsonism and to establish a basis for formulating a public health policy, using an over 10-year follow-up cohort model.

## METHODS

### Statement of Ethics

This research project was approved by the NHIS of South Korea (NHIS-2017-2-542). This study was approved by the Institutional Review Board of our hospital and adhered to the tenets of the Declaration of Helsinki. The need for obtaining informed consent was waived. All authors contributed to the study design, results interpretation, and the decision to submit the manuscript for publication. No commercial support was obtained for this study.

### Database

The management of the NHIS is divided into two independent institutions: the NHIC and Health Insurance Review and Assessment Service of Korea (HIRA). The NHIC has accumulated data (National Health Insurance Database; NHID) of the insured person, premium imposition, and regular health check-up, while HIRA has accumulated data on health insurance claims, which are accompanied by data regarding diagnostic codes, procedures, prescribed medication, personal information, information about the hospital, the direct medical costs of both inpatient and outpatient care, and dental services.^14^ Therefore, we utilized the combined data from the NHIC database of regular health check-ups in 2003 and 2004, and the HIRA database of reimbursement claims from 2003 to 2013.

Blue-collar workers can undergo annual health check-ups, while white-collar workers and self-employed individuals, such as independent businessmen, farmers, fishermen, housewives, and retirees, can do so every other year. The following are regular health check-up items:

1. Body index measurements: height, body weight, waist circumference, body mass index (BMI).
2. History: smoking, alcohol use, and medication history. Medication history includes the drugs used for hypertension, diabetes, and hyperlipidemia.
3. Screening test: visual, hearing, oral hygiene, and laboratory tests (total cholesterol, aspartate aminotransferase, alanine aminotransferase, gamma-guanosine triphosphate, fasting blood glucose, urine protein, plasma creatinine, hemoglobin), and chest X-rays were performed.

After the first health check-up, patients suspected of hypertension or diabetes mellitus (DM) undergo a second health check-up.

### Study Sample

NHIS and HIRA use the Korean Classification of Diseases (KCD) as disease classification codes; these are modified from the International Classification of Diseases. Of the 5,147,950 people who underwent regular health check-ups from 2002 to 2003, 10% were randomly selected (514,795 subjects) for inclusion in this study.

Based on HIRA data, we defined patients with Parkinsonism as those who had KCD codes for Parkinsonism or Parkinson disease (not-otherwise-specified, idiopathic, primary) (G20). We excluded the codes for secondary Parkinsonism (G21) and Parkinsonism in disease classified elsewhere (G22).

### Definition of Risk factors

We defined the following as risk factors: age, hypertension, DM, alcohol drinking, smoking, socioeconomic status, statin medication use, depression, and anxiety. Risk factors were assigned according to the presence of codes in health insurance claims: hypertension (I10 to I15), DM (E10 to E14), depression (F32, F33), and anxiety (F40 or F41). Statin medication use was defined by drug codes for statins in the health insurance claims. Based on the Asian standard, BMI was classified into five grades, i.e., < 18.5 as underweight, 18.5–22.9 kg/m^2^ as normal, 22.9–24.9 kg/m^2^ as overweight, 25–29.9 kg/m^2^ as moderately obese, and 30–35 kg/m^2^ severely obese. Smoking history was classified into non-smokers, ex-smokers, current smokers. Drinking history was divided into the following five grades: no drinking, drinking 1–2 times a week, 2–3 times a week, 3–4 times a week, drinking virtually daily.

### Classification of socioeconomic status by health insurance premium rate

We assumed that the health insurance premium rate reflects the socioeconomic status of subjects. The health insurance premieum payment was divided into three types: “self-employed insured,” “employed insured,” and Medicaid. Medicaid provides medical care for old or disabled individuals who have little income or property. The population on Medicaid pays a part or none of their medical bills, although there are regulations and legal limits to the use of the medical system.

The premium rate of employees is set by a standard that is based on their monthly salary, but as the income of self-employed individuals could not be determined exactly, their premium rate is set according to conversion points that include the insurance holder’s property, such the cost of their house, whether they possess a car, economic activity by age and sex, and total income.

The health insurance premium rate is divided into 10 quantiles for each type of premium payment. We divided socioeconomic status into nine groups according to the premium payment system (self-employed, employed, Medicaid) and premium rate (four grades) (Table 1).

**Table 1.** Classification of socioeconomic class by health insurance premium standard (mean values from 2002 to 2003)

| Classification | Income | Employed Insured | Self-employed Insured |
| --- | --- | --- | --- |
|  | Quantile | KRW (US Dollar) | KRW (US Dollar) |
| Group 1 | 10 | 843,945 (703.29) | 982,721 (818.93) |
| Group 2 | 9 | 569,525 (474.60) | 590,686 (492.23) |
|  | 8 | 384,443 (320.36) | 366,403 (305.34) |
|  | 7 | 259,505 (216.25) | 227,161 (189.30) |
| Group 3 | 6 | 182,225 (151.85) | 140,893 (117.41) |
|  | 5 | 128,145 (106.79) | 93,892 (78.24) |
|  | 4 | 83,720 (69.77) | 64,603 (53.84) |
| Group 4 | 3 | 49,345 (41.12) | 40,541 (33.78) |
|  | 2 | 24,625 (20.52) | 29,120 (247.27) |
|  | 1 | 9,945 (8.29) | 8,531 (7.11) |

\*1\$ = 1200 Korean Won Rate

### Data processing and statistical analysis

We created a 10-year follow-up cohort model for random sampling of 10% of all subjects enrolled in regular health check-ups in 2002 and 2003 (Figure 1). We defined Parkinsonism onset by the presence of a Parkinson’s disease diagnostic code (G20) for the main or second disease in health insurance claims.

**Figure 1.**
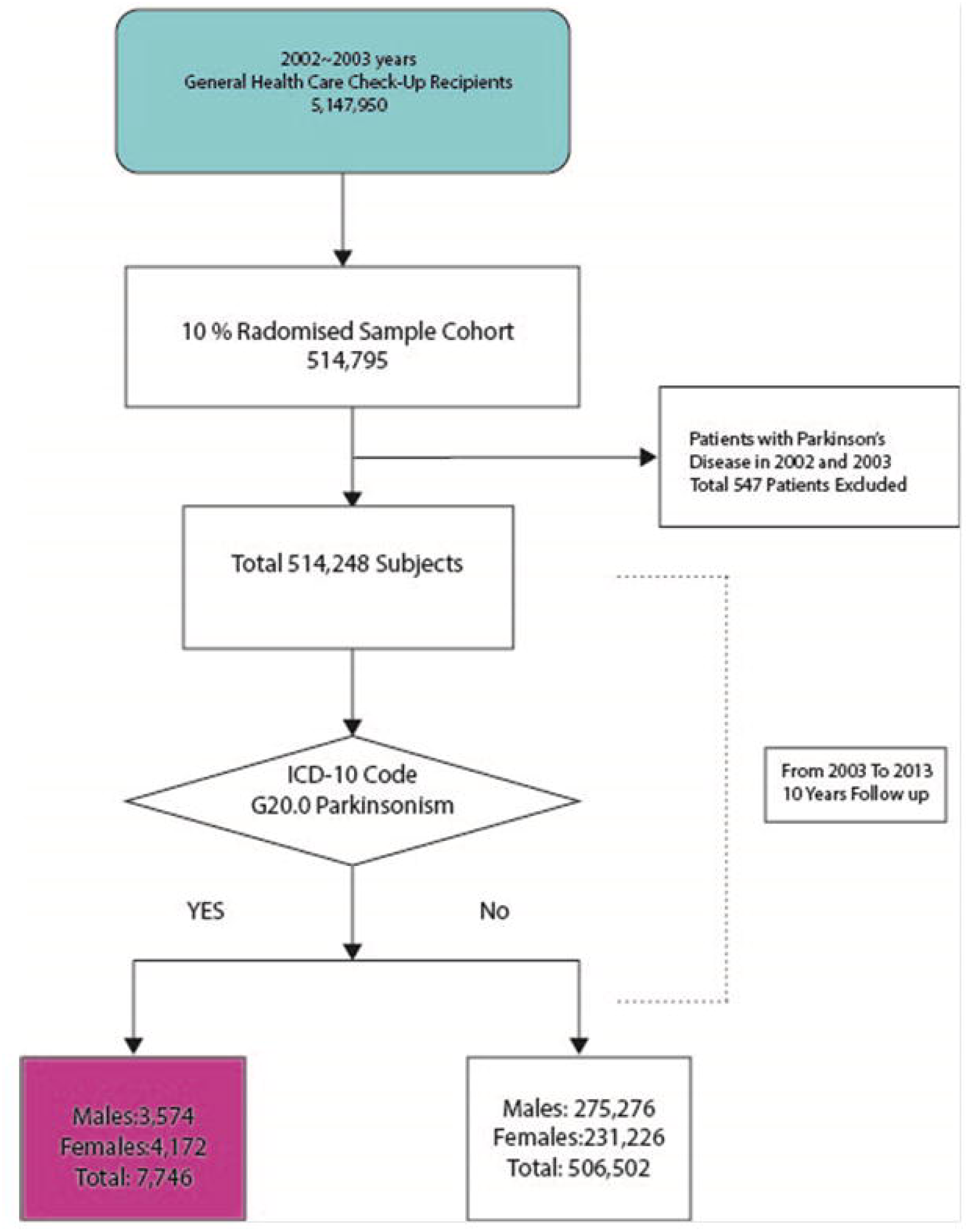
Flowchart of the study design

Descriptive statistics of the study populations are presented, and chi-square tests were performed to examine the association of risk factors with Parkinsonism. Adjusted hazard ratios (HRs) and 95% confidence intervals (CI) were calculated using multivariate Cox proportional hazard regression. A significance level of 0.05 was selected. SAS for Windows, version 9.2 (SAS Inc, Cary, NC, USA) was used to perform statistical analyses.

## RESULTS

Table 2 shows the demographic data and the incidence of Parkinsonism in each group among subjects who underwent regular health check-ups in 2002 and 2003. The HRs of the subjects aged 50–59, 60–69, 70–79, and 80 years or older were 3.101, 8.958, 14.709, and 16.797, respectively, and statistically significant *(p <* 0.0001). When comparing the sexes, the HR of females was 0.971, which was statistically insignificant *(p =* 0.3273) (Table 3). In cases with hypertension, DM, depression, and anxiety, the HRs were 1.259, 1.255, 1.554, and 1.808, respectively, *(p <* 0.0001). The HR of the group taking statins was 1.157, which was higher than that of the group not taking statins at the time of diagnosis.

**Table 2.** Demographic data and the incidence of Parkinsonism

| Group | Subjects with<br>Parkinsonism | Subjects without<br>Parkinsonism | Incidence (%) |
| --- | --- | --- | --- |
| **Age (years)** |  |  |  |
| 40-49 | 648 | 224,862 | 0.13 |
| 50-59 | 1,478 | 144,470 | 0.29 |
| 60-69 | 3,447 | 101,583 | 0.67 |
| 70-79 | 2,088 | 34,265 | 0.41 |
| >80 | 85 | 1,322 | 0.02 |
| **Total** | **7,746** | **506,502** | **1.51** |
| **Sex** |  |  |  |
| Men | 3,574 | 275,276 | 0.69 |
| Women | 4,172 | 231,226 | 0.81 |
| **Hypertension** |  |  |  |
| Yes | 3,917 | 132,388 | 0.76 |
| No | 3,829 | 374,114 | 0.74 |
| **Diabetes** |  |  |  |
| Yes | 2,096 | 67,667 | 0.41 |
| No | 5,650 | 438,835 | 1.10 |
| **Depression** |  |  |  |
| Yes | 1,200 | 24,302 | 0.23 |
| No | 5,258 | 399,874 | 1.24 |
| **Anxiety** |  |  |  |
| Yes | 1,863 | 51,437 | 0.36 |
| No | 5,883 | 455,065 | 1.14 |
| **Statin** |  |  |  |
| Yes | 938 | 32,537 | 0.18 |
| No | 6,808 | 473,965 | 1.32 |
| **BMI** |  |  |  |
| < 18.5 | 181 | 9,659 | 0.04 |
| 18.5–22.9 | 2,108 | 149,450 | 0.50 |
| 23.0-24.9 | 1,681 | 115,366 | 0.40 |
| 25.0-29.9 | 2,007 | 133,134 | 0.47 |
| 30.0-39.9 | 188 | 11,463 | 0.04 |
| **Smoking history (number of missing data = 21,740)** |  |  |  |
| Non-Smoker | 5,691 | 324,584 | 1.16 |
| Ex-smoker | 532 | 43,134 | 0.11 |
| Current smoker | 1,217 | 117,350 | 0.25 |
| **Alcohol use history (numbers of missing data = 21,740)** |  |  |  |
| No | 5,393 | 280,322 | 1.07 |
| 2-3 times/month | 735 | 76,330 | 0.15 |
| 1-2 times/week | 703 | 82,950 | 0.14 |
| 3-4 times/week | 344 | 35,314 | 0.07 |
| Daily | 390 | 22,069 | 0.08 |
| **Socioeconomic status** |  |  |  |
| Medicaid | 39 | 1,006 | 0.01 |
| Self-employed insured grade 1 | 1,136 | 38,514 | 0.22 |
| Self-employed insured grade 2 | 1,004 | 51,036 | 0.20 |
| Self-employed insured grade 3 | 1,009 | 70,598 | 0.20 |
| Self-employed insured grade 4 | 448 | 29,787 | 0.09 |
| Employed-insured Grade 1 | 839 | 75,401 | 0.16 |
| Employed-insured Grade 2 | 767 | 65,357 | 0.15 |
| Employed-insured Grade 3 | 1,649 | 113,355 | 0.32 |
| Employed-insured Grade 4 | 855 | 614,448 | 0.17 |

**Table 3.** Hazard ratios of known risk factors of Parkinsonism

| Variables | HR | 95% Hazard Ratio Confidence Limits | P Value |
| --- | --- | --- | --- |
| **Age** |  |  |  |
| 40s | 1 |  |  |
| 50s | 3.101 | 2.808-3.423 | <.0001 |
| 60s | 8.958 | 8.167-9.826 | <.0001 |
| 70s | 14.709 | 13.319-16.244 | <.0001 |
| 80s | 16.797 | 13.112-21.518 | <.0001 |
| **Sex** |  |  |  |
| Male | 1 |  |  |
| Female | 0.971 | 0.914-1.030 | 0.3273 |
| **Comorbidities** |  |  |  |
| Hypertension | 1.259 | 1.194-1.328 | <.0001 |
| DM | 1.255 | 1.186-1.329 | <.0001 |
| Depression | 1.808 | 1.664-1.965 | <.0001 |
| Anxiety | 1.554 | 1.462-1.652 | <.0001 |
| Statin | 1.157 | 1.072-1.250 | 0.0002 |
| **BMI** |  |  |  |
| 18.5-22.9 | 0.933 | 0.808-1.078 | 0.3458 |
| <18.5 | 1.017 | 0.956-1.083 | 0.5906 |
| 23.0-24.9 | 1.005 | 0.946-1.067 | 0.8739 |
| 25.0-29.9 | 0.932 | 0.806-1.078 | 0.3452 |
| 30.0-39.9 | 1.878 | 1.746-2.021 | <.0001 |
| **Smoking History** |  |  |  |
| Non-Smoker | 1 |  |  |
| Ex-smoker | 0.920 | 0.831-1.018 | 0.1062 |
| Current Smoker | 0.920 | 0.854-0.991 | 0.0287 |
| **Alcohol Drinking History** |  |  |  |
| Non-Alcohol Drinker | 1 |  |  |
| Drinking 2-3 times per month | 0.868 | 0.797-0.946 | 0.0012 |
| Drinking 1-2 times per week | 0.810 | 0.740-0.887 | <.0001 |
| Drinking 3-4 times per week | 0.803 | 0.710-0.908 | 0.0005 |
| Drinking- almost everyday | 0.974 | 0.869-1.092 | 0.6530 |
| **Economic status** |  |  |  |
| Medicaid | 1 |  |  |
| Self-employed insured grade 1 | 0.571 | 0.414-0.788 | 0.0007 |
| Self-employed insured grade 2 | 0.584 | 0.422-0.807 | 0.0011 |
| Self-employed insured grade 3 | 0.544 | 0.393-0.751 | 0.0002 |
| Self-employed insured grade 4 | 0.502 | 0.359-0.701 | <.0001 |
| Employed-insured Grade 1 | 0.453 | 0.327-0.627 | <.0001 |
| Employed-insured Grade 2 | 0.518 | 0.374-0.717 | <.0001 |
| Employed-insured Grade 3 | 0.564 | 0.409-0.776 | 0.0004 |
| Employed-insured Grade 4 | 0.553 | 0.400-0.766 | 0.0004 |

The risk of Parkinsonism was higher in the highest BMI group than in the normal weight group *(p <* 0.0001). The HRs of ex-smokers and current smokers were the same (0.920), but was statistically significant only for current smokers. *(p <* 0.0287). The HRs of all alcohol-drinking groups were < 1 and were statistically significant *(p <* 0.0001), except for the daily alcohol-drinking group *(p =* 0.6530). As Medicaid was set as the standard, the HR was < 1 in all groups *(p <* 0.05), indicating that socioeconomic status and Parkinsonism were closely related.

## DISCUSSION

The results of our study revealed that increasing age, hypertension, DM, depression, anxiety, extreme overweight, statin medication use, non-smoking, non-alcohol drinking, and the lowest socioeconomic class were statistically related to the onset of Parkinsonism.

Parkinsonism is the second most neurodegenerative disease, and its incidence has been reported to increase with age in all studies.^4^ In our study, there was no difference in the incidence of Parkinsonism between the sexes, but one review has reported that males have a 1.5- to 2-fold increased risk of developing Parkinsonism than that of females.^4^ However, some large population-based cohort studies on sex and Parkinsonism have been inconsistent. One study reported that men are more prone to Parkinsonism than women,^5^ and another found the opposite,^15^ whereas other studies found no role for sex.^5, 15, 16^ Therefore, a meta-analysis including large population-based cohort studies is necessary to evaluate the incidence of Parkinsonism according to sex.

Hypertension can cause ischemic cerebrovascular lesions that involve dopaminergic or non-dopaminergic subcortical structures. It also can cause hypertensive vasculopathy in basal ganglia, which may injure the dopaminergic cells in the pars compacta and cause Parkinsonism by breaking the neuronal connections between the substantia nigra and putamen, or by decreasing expression of the (β-2, α-4 subunit of the nicotinic acetylcholine receptor that activates the dopaminergic pathways.^17^ However, most studies related to hypertension and the Parkinsonism incidence yielded results contrary to our findings^7, 13, 18, 19^ or showed no relationship between hypertension and Parkinsonism development.^20^ This is considered to be related to the pathophysiology and progress of Parkinsonism. Autonomic nervous system dysfunction is a very common feature in Parkinsonism.^19^ Loss of sympathetic cardiac innervation in Parkinsonism causes changes in cardiovascular physiology, which may precede Parkinsonism diagnosis.^7, 21, 22^ Because the Lewy body pathology involves the dorsal motor nucleus of the vagus nerve,^23^ parasympathetic tone may become dysregulated, leading to orthostatic hypotension and decreased heart rate variability, which are typical of autonomic dysfunction in Parkinsonism.^24^

We found DM to be a risk factor for the development of Parkinsonism. Cross-sectional studies are limited in their ability to prove causality, and have not yielded concordant results. In one study, patients with Parkinsonism were more likely to have DM, but another study showed opposite results.^19, 25^ However, one large 18-year prospective cohort study reported that type 2 DM was a risk factor for Parkinsonism development, consistent with our findings.^26^ Although the relationship between diabetes and Parkinsonism is unclear, animal and in vitro studies have shown that insulin and brain dopaminergic activity are interrelated.^27^ Thus, insulin dysregulation and the change in insulin action are assumed to affect the pathophysiology and clinical symptoms of Parkinsonism. However, hypertension and DM also increase with aging, as does Parkinsonism (excluding childhood type 1 DM and secondary hypertension in the young). Therefore, we assume that hypertension and DM are risk factors for development of Parkinsonism.

Depression and anxiety are well-known, common, non-motor symptoms of Parkinsonism.^10^ Depression is an early marker of Parkinsonism pathogenesis; significant involvement of dopaminergic neurons in the substantia nigra^10, 28^ and anxiety are early symptoms of Parkinsonism and are known to involve noradrenergic and serotonergic neurons in the brainstem.^29^ Previous studies have shown that depression and anxiety are significantly associated with Parkinsonism.^9, 10, 18, 20, 28^ However, except for one cohort study,^30^ these were cross-sectional studies,^9, 10, 18, 20, 28^ which make it difficult to distinguish between risk factors for the onset of Parkinsonism and non-motor symptoms of Parkinsonism. Nevertheless, our study proved that depression and anxiety are risk factors for Parkinsonism.

Cholesterol is abundant in the central nervous system and plays an important protective role against the early development of Parkinsonism. Higher serum cholesterol and serum triglyceride levels have been reported to reduce the risk of Parkinsonism.^31^ Statin is a 3-hydroxy-3-methylglutaryl-coenzyme A reductase inhibitor of cholesterol synthesis.^32^ Similar to our findings, a previous prospective study also demonstrated that statin use significantly increased the risk of Parkinsonism. However, the findings on statin medication and development of Parkinsonism have been inconsistent.^33, 34^ Most of these studies were cross-sectional and retrospective, making it difficult to identify causal relationships, but a recent meta-analysis reported that statin medication was associated with a lower risk of developing Parkinsonism.^35^

Obesity is a well-known risk factor for metabolic and vascular disorders, such as type 2 DM, coronary heart disease, and stroke.^36^ Therefore, obesity is suspected as a risk factor for Parkinsonism; this study showed that severe obesity elevated the risk of Parkinsonism significantly. The underlying mechanism may involve reduced availability of dopamine D2 receptor in the striatum of obese than non-obese individuals.^11^ However, a meta-analysis reported that higher BMI did not increase Parkinsonism risk;^36^ thus the relationship between obesity and the risk of Parkinsonism remains controversial.^37^

Smoking is a powerful risk factor for hypertension, atherosclerosis, and ischemic and hemorrhagic stroke, but is a well-established preventative factor for Parkinsonism, as also found in our study. Non-smokers more frequently developed Parkinsonism than ex-smokers and current smokers.^12, 13, 18, 22, 28, 37^ Smoking increases dopamine activity by reducing MAO-B activity.^12, 28^ Cytochrome P-450 family members are responsible for the metabolism and detoxification of environmental toxins that cause dopaminergic neural damage. Cytochrome P-450 is induced by smoking, due to the polycyclic hydrocarbons, such as benzopyrene, present in cigarette smoke.^12, 38^

In our study, the risk of Parkinsonism was statistically significantly reduced in all alcohol-drinking groups, except for the daily alcohol-drinking group, as compared with the non-alcohol drinking group.^39^ One study reported that alcohol drinking was associated with increased risk of Parkinsonism,^37^ while the results of other studies were similar to ours.^13, 18^ One review article concluded that a weak protective association was more likely to be found between alcohol drinking and Parkinsonism risk in studies at greater risk of selection and recall bias. One study found that the risk of Parkinsonism was elevated in patients with alcohol-abuse disorder,^6^ while our study showed that there was no lowered risk of Parkinsonism in the daily alcohol-drinking group. Therefore, taken together, it is likely that appropriate alcohol consumption is associated with lowered Parkinsonism risk.

One study reported that the incidence of Parkinsonism was not affected by socioeconomic status.^5^ However, we found that the Parkinsonism risk was reduced in all health insurance payment groups, as compared to the Medicaid group. This is probably because the standards for determining socioeconomic status differed in the two studies, and Medicaid subscribers in South Korea are mainly old individuals with no economic capacity.

The strengths of our study are that we used a nationwide 10-year follow-up cohort model with a population > 500,000, and that the data are relatively objective and accurate, based on regular health check-up data and disease diagnostic codes from HIRA. HIRA reviews claims with disease codes to determine whether reimbursements are clinically valid. Thus, HIRA can maintain the quality of health care and provide the standard medical service guidelines for each disease. There is little chance that medical records were duplicated or omitted, because all Korean residents receive a unique identification number at birth, which is used in medical claims. Moreover, the relationship between the onset of Parkinsonism, various comorbidities, body indexes, and various known risk factors could be verified simultaneously in the present study. We also assessed socioeconomic status based on health premium payment methods. As we used a retrospective cohort model, rather than a cross-sectional study, our data are useful for identification of causality.

The study had the following limitations: subjects < 40 years old were excluded, because the subjects reporting for regular health check-up are ≥ 40 years In a general multicenter prospective cohort study, the same diagnostic criteria are used for data collection; however, since the nationwide data cohort model uses a health insurance-related database, it is very likely that the same diagnostic criteria were not applied. Smoking, BMI, and premium payment type are not fixed, but may have changed during the follow-up period. If an employee becomes unemployed or starts his or her business, the payment system will change from Employed-insured to Self-insured, and in extreme cases, if the insurance holder has no earned income or property, the premium payment form may change to Medicaid.

## Conclusion

As previously reported, we found increasing age, depression, anxiety, and non-smoker status to be risk factors for Parkinsonism. However, sex, hypertension, DM, statin medication use, alcohol drinking, and lower socioeconomic status were also identified as risk factors, but have not been reported as such in previous studies, and thus require verification in future studies. Moreover, the known risk factors of Parkinsonism and socioeconomic status based on the NHID has not been reported for South Korea previously. By comparing these findings with the results of studies in other countries, insights into the pathophysiology and epidemiology of Parkinsonism may be gained, and the results may facilitate formulation of a Korean public health policy. To verify additional risk factors of Parkinsonism, it is necessary to perform studies in which NHID data and genetic and environmental factors are combined.

